# Comparative phyloproteomics identifies conserved plasmodesmal proteins

**DOI:** 10.1101/2022.06.01.494363

**Authors:** Matthew G. Johnston, Andrew Breakspear, Sebastian Samwald, Dan Zhang, Diana Papp, Christine Faulkner, Jeroen de Keijzer

## Abstract

Plasmodesmata connect neighbouring plant cells across the cell wall. They are cytosolic bridges, lined by the plasma membrane and traversed by endoplasmic reticulum to connect these cell components between cells and tissues. While plasmodesmata are notoriously difficult to extract, tissue fractionation and proteomic analyses have yielded valuable knowledge of their composition. Most proteomic profiles originate from cell suspension cultures in which simple plasmodesmata dominate and have been exclusively generated from dicotyledonous plant species. Here we have generated two novel proteomes to expand tissue and taxonomic representation of plasmodesmata: one from mature Arabidopsis leaves and one from the moss *Physcomitrium patens*. We have leveraged these and existing data to perform a comparative analysis that, owing to comparing proteomes from an expanded taxonomic tree, allowed us to identify conserved protein families that are associated with plasmodesmata that likely serve as core structural or functional components. Thus, we identified β-1,3-glucanases, C2 lipid-binding proteins and tetraspanins as core plasmodesmal components, with proteins from *P. patens* and Arabidopsis maintaining plasmodesmal association across diverse species. Our approach has not only identified elements of a conserved, core plasmodesmal proteome, but also demonstrated the added power offered by comparative analysis. Conserved plasmodesmal proteins establish a basis upon which ancient plasmodesmal function can be further investigated to determine the essential roles these structures play in multicellular organism physiology in the green lineages.

## Introduction

Plasmodesmata (PD) are membrane-lined connections that traverse the cell wall and interconnect the cytoplasm, plasma membrane and endoplasmic reticulum between plant cells. The direct cytosol-to-cytosol contact enables the sharing of resources and information, underpinning growth, developmental and response processes (Benitez-Alfonso, 2014; Sevilem et al., 2015; Tilsner et al., 2016; Brunkard and Zambryski, 2017; Cheval and Faulkner, 2018; Sager and Lee, 2018). PD are dynamic, adapting their conductivity in response to internal and external cues to create transient domains within tissues. While it is established that their responses to a range of environmental signals are enabled by specialized signaling machinery (Stahl and Faulkner, 2016; Grison et al., 2019; Cheval et al., 2020), we still have little understanding of the molecular machinery that brings about their biogenesis and structure.

To obtain a comprehensive overview of proteins present at PD, and ultimately build understanding of their role in physiology and development, proteomic characterisation of plasmodesmata-enriched fractions has been performed on multiple occasions (Faulkner et al., 2005; Fernandez-Calvino et al., 2011; Leijon et al., 2018; Brault et al., 2019; Park et al., 2017). Indeed, such proteomes have yielded extremely valuable insights into plasmodesmal structure and function, identifying novel plasmodesmal machinery that has been leveraged to gain further understanding of the function and relevance of plasmodesmata in a range of plant processes. To date, these approaches have yielded knowledge of how PD contribute to lateral root formation (Benitez-Alfonso et al., 2013) and immune signalling (Faulkner et al., 2013). In addition to revealing novel plasmodesmal proteins, proteomic analyses generate an expanding ‘parts list’ that allows us to ask whether recurrent protein classes are found at PD in multiple plant tissues and species, and thus define a core protein complement of plasmodesmata (Kirk et al., 2021). However, sampling across differentiated tissues and taxonomic groups in PD proteomic studies is hitherto poor, limiting the scope of such an approach. As PD are understood to be a feature conserved across land plants (Brunkard and Zambryski, 2017), expanding our current knowledge relating to the plasmodesmata of flowering plants to extant species belonging to different taxonomic groups would give greater insight into core and conserved plasmodesmal components.

The bryophytes are a group of plants sister to the vascular plants (tracheophytes), with these clades diverging soon (∼445 Mya) after the conquest of the land by the green kingdom (∼490 Mya) (Morris et al., 2018). Electron microscopy has revealed PD across the tissues of bryophyte share architectural features such as an outer plasma membrane lining and a central desmotubule (comprised of endoplasmic reticulum), with those in flowering plants (Ligrone, 1994; Ligrone and Duckett, 1998; Cook et al., 1997; Ligrone et al., 2000). These observations suggest PD are a trait present in the ancestor of all land plants and that elements of their structure observed across diverse extant species might be essential to plasmodesmal function, being conserved or repeatedly recruited to plasmodesmata. Except for a limited analysis of the proteins present in plasmodesmata of the giant-celled green alga *Chara corallina* (Faulkner et al., 2005), molecular details about the composition of plasmodesmata outside the flowering plants are lacking, leaving questions of the molecular conservation of plasmodesmata unanswered.

A comparison of extant traits and molecular constituents between living ancestors would provide a powerful entry point towards establishing which plasmodesmal components are core, and which are derived. In recent years, extant species from Bryophyta such as *Marchantia polymorpha* and *Physcomitrium* (formerly *Physcomitrella*) *patens* have grown to be important models for plant research, (Rensing, 2017; Rensing et al., 2020; Naramoto et al., 2022; Bowman, 2022). This presents an opportunity to exploit recent developments in methods for extracting plasmodesmata from differentiated green tissues to phylogenetically expand information relating to the molecular composition of plasmodesmata. Here, we have generated new plasmodesmal proteomes from differentiated tissue of *Arabidopsis thaliana* and *Physcomitrium patens*. Leveraging these and existing proteomes we performed a comparative phylogenetic analysis to identify core plasmodesmal proteins. This analysis revealed 16 protein orthogroups that are represented across most proteomes. We analysed five of these orthogroups and validated β-1,3-glucanases, C2 lipid-binding proteins and tetraspanins as containing conserved, core plasmodesmal proteins. Thus, we have demonstrated that comparative proteomics can reveal essential features of plasmodesmata, with the potential to define basic rules and requirements of symplastic cell-to-cell communication in the multicellular green lineage.

## Materials and Methods

### Plant material and growth conditions

For plasmodesmal extraction, *Arabidopsis thaliana* Col-0 plants were grown on soil in short day conditions (10 h light / 14 h dark) at 22°C. Leaves were harvested five weeks after germination. For stable transformation, *A. thaliana* plants were grown in long day conditions (16 h light/8 h dark). *Physcomitrium* (*Physcomitrella*) *patens* tissues for generating plasmodesmal fractions was grown on BCDAT medium in long day conditions (16 h light / 8 h dark) at 25 °C. Protonemal tissue was grown on top of nitrocellulose membrane for 1 week, whereas gametophore tissue was grown directly on the medium for 4 weeks. Routine *P. patens* culture for generating and maintaining transformants was performed under continuous light at 25 °C on BCD-AT medium. *Nicotiana benthamiana* plants were grown on soil with 16 h light / 8 h dark at 23 °C.

### Plasmodesmal purification

Plasmodesmata were extracted from expanded rosette leaves of 5-week-old Arabidopsis plants and a mix of *Physcomitrium patens* protonemal and gametophore tissue. To fully homogenise differentiated tissue, we extracted plasmodesmata according to (Cheval et al., 2020), with the key difference in approach from (Faulkner and Bayer, 2017) being the homogenisation method and the use of Triton X-100 to disrupt chloroplasts. First, frozen mature tissue was ground into a powder in liquid nitrogen and suspended with extraction buffer (EB: 50 mM Tris-HCl pH 7.5, 150 mM NaCl, 1 × cOmplete™ ULTRA protease inhibitors (Sigma), 1 mM PMSF, 1% (w/v) PVP-40kDa (Sigma)) and ultrasonicated for 1 minute in six 10-second pulses with a five second pause between each pulse (Soniprep 150 Plus, MSE). The sample was passed twice through a high-pressure homogenizer (EmulsiFlex™-B15, Avestin) at 80 PSI. Triton X-100 (10% v/v) was added dropwise to the resultant homogenate to a final concentration of 0.5% (v/v) to disrupt residual chloroplasts and cell walls were collected by centrifugation at 400*g*. The cell wall pellet was washed three times (four for *P. patens* samples) with EB (15 mL) and centrifuged at 400*g*. We validated the method for *P. patens* by calcofluor staining of cell walls at the different stages of fractionation (Fig S1) showing that the size of cell wall fragments generated by this approach are similar to those derived from *A. thaliana* suspension cells (30 – 100 µm) (Grison et al., 2015).

The cleaned cell wall pellet was incubated in an equal volume of cellulase buffer (CB: 20 mM MES-KOH pH 5.5, 100 mM NaCl, 2% w/v Cellulase R-10 (Yakult Pharmaceutical Co., Ltd., Japan), 1 × cOmplete™ ULTRA protease inhibitors (Sigma), 1 mM PMSF) for 1 h at 37°C, 200 rpm. Undigested cell wall was removed by centrifugation at 5000*g*, and the supernatant collected as the plasmodesmal membrane containing fraction. The cell wall pellet was washed again with CB to extract residual plasmodesmal membranes and the soluble fraction was ultracentrifuged at 135,000*g* for 1 h. The membrane pellet was resuspended in 50 mM Tris-HCl pH 7.5, 150 mM NaCl, 5 mM DTT, 1×cOmplete™ ULTRA EDTA-free protease inhibitors (Sigma), 1 mM PMSF, 0.2% (v/v) IPEGAL®CA-630 (Sigma).

### Mass spectrometry

Plasmodesmal samples were run 5 mm into a 1.5 mm thick 10% polyacrylamide TRIS resolving gel (containing 0.1% SDS) without a stacking gel, in a glycine 0.1% SDS running buffer. The gel was washed in dH_2_O and then the band was excised. The bands were washed four times in 20% acetonitrile at 40°C for 15 minutes to remove detergents, and then stored at 4°C with 100 µL of dH_2_O.

Mass spectrometry analysis was performed by the Cambridge Centre of Proteomics. 1D gel bands were cut into 1 mm^2^ pieces, destained, reduced (DTT) and alkylated (iodoacetamide) and subjected to enzymatic digestion with trypsin overnight at 37°C. Digested peptides were analysed by LC-MS/MS with a Dionex Ultimate 3000 RSLC nanoUPLC (Thermo Fisher Scientific Inc, Waltham, MA, USA) system and a Q Exactive Orbitrap mass spectrometer (Thermo Fisher Scientific Inc, Waltham, MA, USA). Separation of peptides was performed by reverse-phase chromatography at a flow rate of 300 nL/min and a Thermo Scientific reverse-phase nano-Easy-spray column (Thermo Scientific PepMap C18, 2 µm particle size, 100A pore size, 75 µm i.d. x 50 cm length). Peptides were loaded onto a pre-column (Thermo Scientific PepMap 100 C18, 5 µm particle size, 100 A pore size, 300 µm i.d. x 5 mm length) from the Ultimate 3000 autosampler with 0.1% formic acid for 3 minutes at a flow rate of 15 µL/min. After this period, the column valve was switched to allow elution of peptides from the pre-column onto the analytical column. Solvent A was water + 0.1% formic acid and solvent B was 80% acetonitrile, 20% water + 0.1% formic acid. The linear gradient employed was 2-40% B in 90 minutes (the total run time including column washing and re-equilibration was 120 minutes).

The LC eluant was sprayed into the mass spectrometer by means of an Easy-spray source (Thermo Fisher Scientific Inc.). All m/z values of eluting ions were measured in an Orbitrap mass analyzer, set at a resolution of 70,000 and scanned between m/z 380 - 1,500 Data dependent scans (Top 20) were employed to automatically isolate and generate fragment ions by higher energy collisional dissociation (HCD, Normalised collision energy (NCE): 25%) in the HCD collision cell and measurement of the resulting fragment ions was performed in the Orbitrap analyser, set at a resolution of 17,500. Singly charged ions and ions with unassigned charge states were excluded from being selected for MS/MS and a dynamic exclusion of 20 seconds was employed.

Post-run, all MS/MS data were converted to mgf files and the files were then submitted to the Mascot search algorithm (Matrix Science, London UK, version 2.6.0) and searched against the Cambridge Centre of Proteomics database, including common contaminant sequences containing non-specific proteins such as keratins and trypsin. Variable modifications of oxidation (M) and deamidation (NQ) were applied, as well as a fixed modification of carbamidomethyl (C). The peptide and fragment mass tolerances were set to 20 ppm and 0.1 Da, respectively. A significance threshold value of p < 0.05 and a peptide cut-off score of 20 were also applied. All data (DAT files) were then imported into the Scaffold program (Version 4.10.0, Proteome Software Inc, Portland, OR). Proteins were classed as positively identified when the peptide and protein identification probability thresholds were greater than 95% (Leijon et al., 2018) and proteins were identified in at least two replicates.

### GO Analysis

Gene ontology (GO) was used to test gene lists for cellular localisation enrichment (Carbon et al., 2019; Ashburner et al., 2000). Cellular localisation GO term overrepresentation test was performed, using the Panther database (release 28/07/2020) (Thomas et al., 2006, 2003; Mi et al., 2019) and GO Ontology database (released 10/09/2020) with a Fisher’s exact test and FDR reported. *P. patens* genes were annotated bioinformatically using phylogenetic backpropagation of GO terms via the Panther database (Gaudet et al., 2011). Graphs were drawn using ggplot2 in R (v4.0.0) (Wickham, 2011).

### Bioinformatic analysis

HMMER v3.3 was used for sequence similarity searches (Eddy, 1998). The *P. patens* plasmodesmal proteome was downloaded as peptide sequences from UniProt and used as the reference database for a ‘phmmer’ search against which the *A. thaliana* UniProt proteome was run (UP000006548, accessed 24/04/2020) (Cheng et al., 2017). Protein matches were filtered at either E < 1 × 10^−100^ or E < 1 × 10^−50^ as stated in the text.

Orthofinder (v2.2.6) was used to create *de novo* orthogroups (Emms and Kelly, 2019, 2015). Plasmodesmal proteome protein sequences were downloaded using UniProt, TAIR (Araport11), and Phytozome v12.1 (*Populus trichocarpa* v3.1). Orthofinder was run on these sequences with default settings.

### Phylogenetic analysis

A peptide sequence was downloaded from UniProt for each protein within an orthogroup. The protein FASTA sequences were aligned with Clustal Omega (v1.2.4, (Sievers et al., 2011; Madeira et al., 2019)) to build a consensus sequence. The consensus sequence, in Stockholm format, was used as the basis for a hmmsearch (EBI, HmmerWeb version 2.41.1, (Potter et al., 2018)). A search was conducted against the EMBL Reference Proteomes database restricted to *A. thaliana* (taxon id: 3702), *P. patens* (taxon id: 3218), and *P. trichocarpa* (taxon id: 3694) species sequences with a sequence E-value cut off of 1 × 10^−100^, unless otherwise stated. Protein sequences were manually deduplicated for each gene.

The FASTA sequences for all identified homologues, from the hmmsearch, in all three species were downloaded and a bootstrapped non-rooted phylogenetic was generated using the ‘standard_trimmed_phyml_bootstrap’ ete workflow (v3.1.1, (Huerta-Cepas et al., 2016)). In this workflow, sequences are aligned with Clustal Omega, trimmed with TrimAI (Capella-Gutierrez et al., 2009), and a phylogeny determined with 100 bootstraps using PhyML (Guindon et al., 2010). Trees were drawn using ggtree in R (v4.0.0) (Yu et al., 2017).

### Construct generation for protein tagging in moss

For mNeonGreen tagging of moss candidate plasmodesmal proteins first a mNeonGreen tagging vector was generated. For this the mNeonGreen coding sequence was amplified using primers mNG-HindIII-F and mNG-stop-EcoRI-R (Table S6) from pPY22 (Addgene plasmid #137082; Yi and Goshima, 2020), introducing a GSGGSG-encoding linker before mNeonGreen in the process). Next, using *Hind*III and *Eco*RI restriction sites the citrine fluorophore in pCTRN-nptII (Hiwatashi et al., 2008) was exchanged with the amplified mNeonGreen encoding sequence, resulting in plasmid pmNG-nptII.

Moss mNeonGreen in locus tagging constructs were assembled using InFusion recombination of PCR-amplified fragments. Four fragments were assembled: A vector backbone sequence amplified from pmNG-nptII using primers pBS-vec-PmeI-F and pBS-vec-PmeI-R, two gDNA-amplified homology-arms of approximately 1 kb in length located upstream and downstream of the intended mNeonGreen integration site and a mNeonGreen encoding fragment, which in case of C-terminal fusions, was followed by a G418-resistance cassette (both amplified from pmNG-nptII using primers mNG-noStart-F + mNG-noStart-R or Link-mNG-F + Cassette-R respectively). The resultant plasmids were verified by sequencing and linearized by *Pme*I digestion prior to transformation into *P. patens*.

### Construct generation for N. benthamiana / A. thaliana expression

For localization analysis of putative plasmodesmal candidates by expression in *N. benthamiana* and *A. thaliana* tissues, binary vectors containing the coding sequence of the protein of interest fused to a fluorescent protein were generated. Typically, the coding sequence of a gene of interest was synthesized (Genewiz, China) as a Golden Gate L0 module (Engler et al., 2014) in a pUC57 backbone, except for moss tetraspanin A9TQE7 (Pp3c4_3550V3), moss β-1,3-glucanase A0A2K1J8R8 (Pp3c16_15860V3), and Arabidopsis β-1,3-glucanase Q9FHX5 (AT5G42100; BG_PPAP) (see below). For synthesis, internal *Bsa*I and *Bpi*I restriction sites were removed via silent mutation and appropriate 4-bp overhangs were added to enable Golden Gate cloning (See table S7). Via a *Bsa*I-mediated level 1 Golden Gate reaction coding sequences were linked to an eGFP- or mRuby-encoding fragment with 35S promoter and terminator regions placed upstream and downstream respectively. Coding sequences for moss tetraspanin A9TQE7 and β-1,3-glucanase A0A2K1J8R8 were amplified from moss cDNA, and assembled into a Golden Gate level 0 acceptor plasmid, removing internal *Bsa*I and *Bpi*I sites in the process. For the β-1,3-glucanase, during fragment assembly an mNeonGreen coding fragment was fused in frame after the sequence encoding for the catalytic domain. The tagging construct for Q9FHX5 (BG_PPAP) was generated by inserting Citrine downstream of its predicted signal peptide and assembling the fusion in a Level 1 binary vector with a 35S promoter and terminator.

### Plant transformation

*Arabidopsis thaliana* was transformed by floral dip (Clough and Bent, 1998). *N. benthamiana* was transformed by co-infiltration of *Agrobacterium tumefaciens* (GV3101 (pMP90)) strains harbouring either a binary plasmid encoding for *in planta* expression of the transgene of interest or the p19 silencing suppressor. Leaves were imaged 2 days post infiltration.

*P. patens* was transformed using PEG mediated protoplast transformation (Nishiyama et al., 2000). For constructs without resistance cassette (i.e., these used for N-terminal or internal tagging of the protein of interest), the plasmid p35S-LoxP-HygR (pTN186; Genbank AB542059.1) was co-transformed in a 1:1 ratio such that a first selection step on Hygromycin B containing medium could be performed. Transformants with the correct single integration of the mNeonGreen expression constructs were identified using PCR.

### Confocal imaging

*N. benthamiana* and *A. thaliana* leaf tissue was cut into 1 cm^2^ samples and mounted adaxially. Samples were imaged on a ZEISS LSM800 confocal microscope with a 63×/1.2 water immersion objective lens (C-APOCHROMAT 63×/1.2 water). GFP and mCitrine were excited at 488 nm with an argon laser and collected at 500 – 545 nm. mRuby was excited at 561 nm with a DPSS laser and collected at 590 – 620 nm. Aniline blue (0.1% w/v in 1× PBS pH 7.4) was infiltrated adaxially and excited at 405 nm with a UV laser and collected at 430 – 470 nm. Wall fractions were stained with 0.01% Calcofluor white M2R (F3543, Sigma) and imaged by confocal microscopy with a 20× objective (PLAN APOCHROMAT NA 0.8). Calcofluor white was excited at 405 nm with a UV laser and collected at 430 – 470 nm.

Moss protonemal cells were observed using a 39 mm diameter glass bottom dish, prepared with solidified BCD medium and grown for 4-6 days in a thin layer of the same medium except solidified with 0.7% (w/v) low melting agarose. For all moss fluorescence microscopy experiments the second and third caulonemal cells relative to the tip of a protonemal filament were used. Imaging of endogenous moss proteins tagged with mNeonGreen was performed on a spinning disk confocal microscope consisting of a Nikon Ti-eclipse body equipped with a Yokogawa CSU-X1 spinning disk head and 100× Plan Apo VC objective (NA 1.40). Image digitization was performed with a Photometrics Prime 95B cMOS camera with a 1.2x post-magnification fitted in front of the camera. Typical exposures used were 500-3000ms. For excitation of mNeonGreen a 491 nm laser line was used and emitted light was filtered using a 527/60 bandpass emission filter. All microscope components were operated by MetaMorph software. Colocalization of aniline blue-stained callose deposits with mNeonGreen-tagged proteins of interest was performed on a Leica Stellaris 5 confocal microscope. Aniline blue prepared as a 1.6% (w/v) stock solution in 0.1 M phosphate buffer (pH 8.5) according to (Muller et al., 2022) was diluted in water to a final concentration of 0.02% and then added to the imaging dishes for 48 hrs prior to observation (except for co-localization of beta-1,3-glucanase A0A2K1J8R8 / Pp3c16_15860 where a 2 hr incubation was used). Cells were imaged using a 100× HC plan apo objective (NA 1.40). Excitation of aniline blue was done using a 405 nm solid state laser and emitted light was collected between 420 and 490 nm on a HyD detector with the pinhole set to 0.6 Airy units. Excitation of mNeonGreen was done using 505 nm laser light obtained from a pulsed white light laser and emitted light was collected between 515 and 560 nm on a HyD detector, with the pinhole aperture set to 1 Airy unit. Frames of the two different probes were collected successively and a line-averaging factor of 8 was used.

## Results

### Generation of a plasmodesmal proteome from mature Arabidopsis leaves

There are currently two published plasmodesmata proteomes of *Arabidopsis thaliana* (Brault et al., 2019; Fernandez-Calvino et al., 2011). Both studies followed very similar extraction protocols, generating proteomes from suspension culture cells, followed by differing mass spectrometry techniques. To define a novel *Arabidopsis thaliana* plasmodesmata proteome that represented differentiated tissue, we extracted plasmodesmata from expanded leaves of mature 5-week-old plants and characterised the proteome of the rosette leaves by mass spectrometry. Proteins were considered positively identified in the same manner as (Leijon et al., 2018); if the protein (95% certainty; Searle, 2010) was represented in at least two of the three samples by at least one peptide (95% certainty; Keller et al., 2002). With these thresholds, 238 proteins were identified in the fraction (Table S1).

To assess if the mature leaf plasmodesmal fraction has sufficient purity to define a plasmodesmal proteome, we assessed the cellular localisation GO enrichment of different proteomes. The mature leaf proteome was benchmarked against the existing Arabidopsis cell culture proteomes, noting that the Fernandez-Calvino et al. (2011) proteome defines the plasmodesmal GO term. For ease of referencing, hereafter the proteomes are named ‘AtC1’ (Fernandez-Calvino et al., 2011), ‘AtC2’ (Brault et al., 2019) and ‘AtL’ the proteome from mature leaves produced in this study. All three proteomes were significantly enriched for plasmodesmata labelled proteins (Fig 1). Moreover, all proteomes were significantly enriched for “cell wall” and “plasma membrane” proteins. As plasmodesmata have both cell wall and plasma membrane components, these categories likely contain both potentially undiscovered plasmodesmal proteins and contaminants. The enrichment factor filtering used to define the AtC2 proteome (Brault et al., 2019) worked extremely well, with other likely contaminant categories (e.g., “Golgi apparatus” or “chloroplast”) not over-represented, unlike the unfiltered proteomes. However, the similarity between the representation of proteins in non-plasmodesmal cell components in AtC1 and AtL suggests that that latter is of comparable quality and defines a list of candidate plasmodesmal proteins from Arabidopsis leaves.

**Figure 1.**
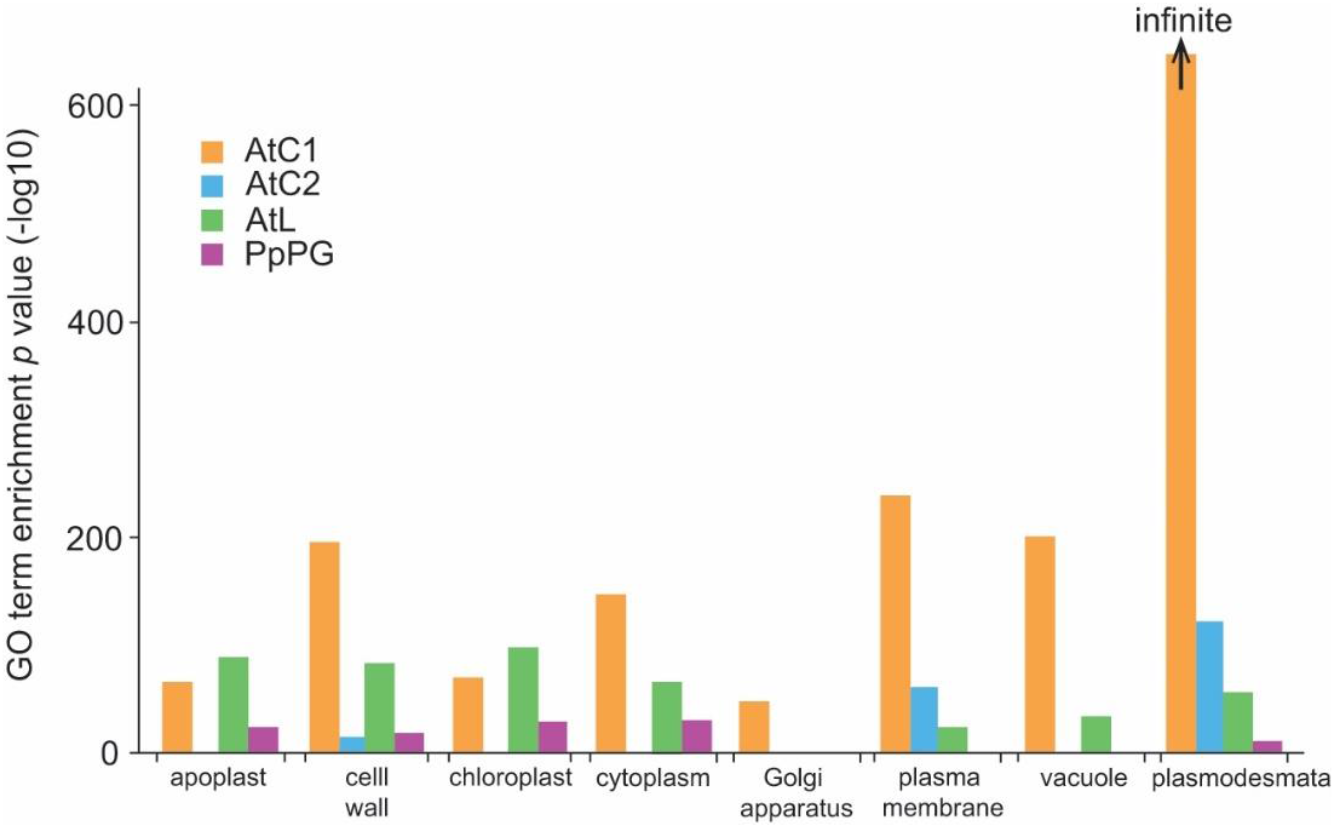
The Arabidopsis plasmodesmal fraction derived from expanded rosette leaves and the moss plasmodesmal fraction derived from protonema and gametophore tisssue are enriched in plasmodesmal proteins. *p*-values for cell compartment GO term enrichment of plasmodesmal proteomes from Arabidopsis cell suspension cultures (AtC1 and AtC2), expanded rosette leaves (AtL), and moss protonema and gametophore tissue (PpPG).

### Generation of a plasmodesmal proteome from P. patens

In addition to Arabidopsis, a plasmodesmal proteome from *Populus trichocarpa* suspension culture cells exists (Leijon et al., 2018), but none are available beyond dicotyledonous flowering plants. To expand the phylogenetic representation of plasmodesmal proteomes, we defined a novel plasmodesmal proteome from the moss *Physcomitrium patens* (termed ‘PpPG’). We extracted plasmodesmata from a mix of protonema and gametophore tissue and analysed the fraction by mass spectrometry. Proteins were identified in the same manner as for the Arabidopsis leaf proteome generating a list of 215 candidate plasmodesmal proteins (Table S2). We confirmed this extraction protocol works in *P. patens* by checking for enrichment of proteins annotated with the plasmodesmal GO term. 185 (86%) of the UniProt identifiers were mapped to the GO Ontology database, with plasmodesma-annotated proteins over-represented (7 proteins, *p* = 3.19 × 10^−5^, 0.51 proteins expected) in the *P. patens* plasmodesmal fraction (Fig 1). The values are reduced due to poor annotation of *P. patens* proteins within the plasmodesmata GO ontology via phylogenetic backpropagation of Arabidopsis GO terms (Gaudet et al., 2011). Nonetheless, we concluded that the employed plasmodesmata extraction protocol produces a plasmodesmata enriched fraction in *P. patens*.

### Phylogenetic comparison of Arabidopsis, poplar and moss plasmodesmal proteomes reveals orthogroups containing core proteins

To further characterise and compare the composition of the *P. patens* plasmodesmal proteome we explored different bioinformatic approaches to find orthologous proteins. First, we used a one-to-one homologue database search approach. Using InParanoid 8.0 (pairwise BLAST, defining orthogroups from an ancestral protein sequence) and MetaPhOrs (defining orthogroups from a meta-analysis of many homologue databases) we converted the *P. patens* protein identifiers to their *A. thaliana* homologue identifier (Table S3). Only 62 (InParanoid) and 52 (MetaPhOrs) *P. patens* proteins were matched to Arabidopsis proteins by this approach, but performing a GO term analysis on these two lists of Arabidopsis identifiers revealed enrichment of the Plasmodesmata GO term (*p* = 7.11 × 10^−16^ and 2.82 × 10^−8^ for InParanoid and MetaPhOrs respectively). However, the low percentage of *P. patens* protein homologues identified (29 and 24%) by this method is too low to allow for the *P. patens* proteome to offer significant power in a comparative analysis.

Our next analysis involved comparing one-to-many, instead of relying on databases to convert *P. patens* proteins to *A. thaliana* homologues. To this end, we used HMMER (v3.3, profile hidden Markov models) to find the closest homologue for *P. patens* plasmodesmal proteins in *A. thaliana*. Using two arbitrary thresholds of E < 1 × 10^−50^ and E < 1 × 10^−100^, HMMER matched 147 (68%) and 80 (37%) *P. patens* proteins to *A. thaliana* proteins, respectively. Even at these conservative values, a HMMER search matched more proteins than database lookup tools. However, one-to-many mapping makes it difficult to translate the *P. patens* proteome members to specific *A. thaliana* proteins. One approach would be to take the most significant (i.e., most likely) homologue for each protein. However, taking *P. patens* A0A2K1JXU2 (“X8 domain-containing protein”; Associated locus Pp3c10_5480V3) as an example, there are two almost indistinguishable top hits in *A. thaliana*: O49737 (E = 4.2 × 10^−101^) and Q8L837 (E = 6.3 × 10^−101^), suggesting it is likely the ancestral protein of A0A2K1JXU2 has undergone a duplication event in *A. thaliana* giving two equally likely homologues. In essence, this builds orthogroups restricted to one *P. patens* member.

Another consideration when using HMMER to assign homologues is that to find phylogenetically conserved proteins, *i*.*e*., to concurrently compare several lists among several species, one list would have to be chosen as the reference frame. Defining the *P. patens* proteome as the reference list allows the distribution of *P. patens* hits across the Arabidopsis proteomes to be compared, but any nuance from comparison between the *A. thaliana* proteomes is lost. Therefore, we tried a third, many-to-many approach by forming *de novo* orthogroups using the OrthoFinder software (Emms and Kelly, 2019).

OrthoFinder uses a pairwise BLAST approach to build orthogroups from an input set of protein sequences. We used OrthoFinder (v2.2.6) to define orthogroups (OGs) between five plasmodesmal proteomes: AtC1, AtC2, AtL, PpPG and the *Populus trichocarpa* cell suspension culture proteome (‘PtC’, Leijon et al., 2018). This analysis returned 992 orthogroups, of which 289 had more than one member and 288 contained proteins from multiple proteomes. Two orthogroups had members from all proteomes, and 17 had members from four of the five proteomes (Table 1, Table S4). We noted that members of the IMK2 OG (OG50) are receptor-like kinases belonging to the LRR III group represented by OG9. Similarly, the sole member of the calcium-dependent lipid-binding orthogroup identified in the Arabidopsis proteomes shows similarity to members of the C2 lipid-binding orthogroup (OG3, phmmer search E= 9.6 × 10^−6^). Therefore, OG50 and OG3 were not considered independently. Further, while OG19, representing DUF26 containing proteins that include the PDLPs, is represented in the proteomes from *P. trichocarpa* and *A. thaliana*, it does not have any *P. patens* homologues (Vaattovaara et al., 2019) and so we excluded it as a candidate core orthogroup. We defined the remaining 16 orthogroups as containing proteins that are ‘phylogenetically conserved plasmodesmal proteins’ (Table 1).

**Table 1.** List of orthogroups identified in at least four of five proteomes

| Orthogroup | Protein Class | # Proteomes | # Proteins | Localised at PD |
| --- | --- | --- | --- | --- |
| OG0 | β-1,3-glucanase | 5 | 27 | (Levy et al., 2007; Benítez-Alfonso et al., 2013) |
| OG1 | Peroxidase | 4 | 22 | No |
| OG3 | C2 lipid-binding | 4 | 16 | (Brault et al., 2019) |
| OG4 | SKU5 | 4 | 13 | No |
| OG5 | GDSL esterase/lipase | 4 | 13 | No |
| OG6 | Tetraspanin | 4 | 12 | (Fernandez-Calvino et al., 2011) |
| OG7 | ATP-binding cassette | 4 | 11 | No |
| OG8 | Aspartyl protease | 4 | 10 | No |
| OG9 | Leucine-rich repeat receptor-like kinase | 4 | 10 | (Fernandez-Calvino et al., 2011; Grison et al., 2019) |
| OG10 | Leucine-rich repeat extensin-like | 4 | 10 | No |
| OG13 | Histone H2B | 4 | 9 | No |
| OG14 | Tubulin beta-7 | 4 | 9 | (Blackman and Overall, 1998) |
| OG16 | RNA-binding glycine-rich protein | 4 | 8 | No |
| OG18 | Inflorescence meristem receptor-like kinase 2 | 5 | 7 | No |
| OG19 | DUF26 containing protein | 4 | 7 | (Thomas et al., 2008) |
| OG28 | Eukaryotic translation initiation factor 4A | 4 | 6 | No |
| OG40 | Subtilisin-like protease | 4 | 5 | No |
| OG50 | Calcium-dependent lipid-binding | 4 | 4 | No |
| OG63 | Ribosomal protein | 4 | 4 | No |

### Core moss orthogroup members are plasmodesmal proteins

Rationalising that plasmodesmata are defined by specialised membranes, we first considered orthogroups for which the representatives detected in the Arabidopsis proteomes have at least *in silico* support for membrane association (i.e., either predicted transmembrane helices or an omega site for GPI-anchor attachment). This led us to refine our initial OGs of interest to: OG0 (β -1,3-glucanase), OG3 (C2 lipid-binding), OG6 (Tetraspanin), OG7 (ATP-binding cassette), and OG9 (LRR RLK III). Proteins from OG0 (Levy et al., 2007; Rinne et al., 2011; Benitez-Alfonso et al., 2013), OG3 (Brault et al., 2019), OG6 (Fernandez-Calvino et al., 2011) and OG9 (Fernandez-Calvino et al., 2011; Grison et al., 2019) have already been validated as plasmodesmata-associated by live imaging of fluorescent protein fusions. We selected OG0, OG3, OG6 and OG7 and identified *P. patens* homologues, all bar one present in the *P. patens* plasmodesmal fraction, and further characterized their *in vivo* localization in the native tissues.

For OG0, three *P. patens* β-1,3-glucanases were present in the plasmodesmal fraction (Tables S2, S4). We noted that the protein A0A2K1K5L9 (associated locus Pp3c8_940V3) had an incomplete catalytic domain, and therefore disregarded it for localization analysis. We selected A0A2K1J8R8 (Pp3c16_15860V3), a β-1,3-glucanase with a predicted GPI-anchor similar to most known plasmodesmata-associated β-1,3-glucanases (Levy et al., 2007; Gaudioso-Pedraza and Benitez-Alfonso, 2014; Zavaliev et al., 2016), as a moss representative of OG0. We generated a transgenic *P. patens* line that expresses a fluorescent protein fusion by inserting a mNeonGreen (mNG) encoding sequence at the native genomic locus downstream of the predicted catalytic domain and before the predicted omega site for GPI anchor attachment (Fig S2). Live imaging of *P. patens* protonema shows A0A2K1J8R8-mNG has a punctate localisation at the cell junctions (Fig 2A). Co-localisation with aniline blue (Tomoi et al., 2020; Muller et al., 2022), suggests this fluorescence pattern is co-incident with staining of plasmodesmal callose (Fig 2E) and therefore that A0A2K1J8R8 is a plasmodesmata-associated β-1,3-glucanase.

**Figure 2.**
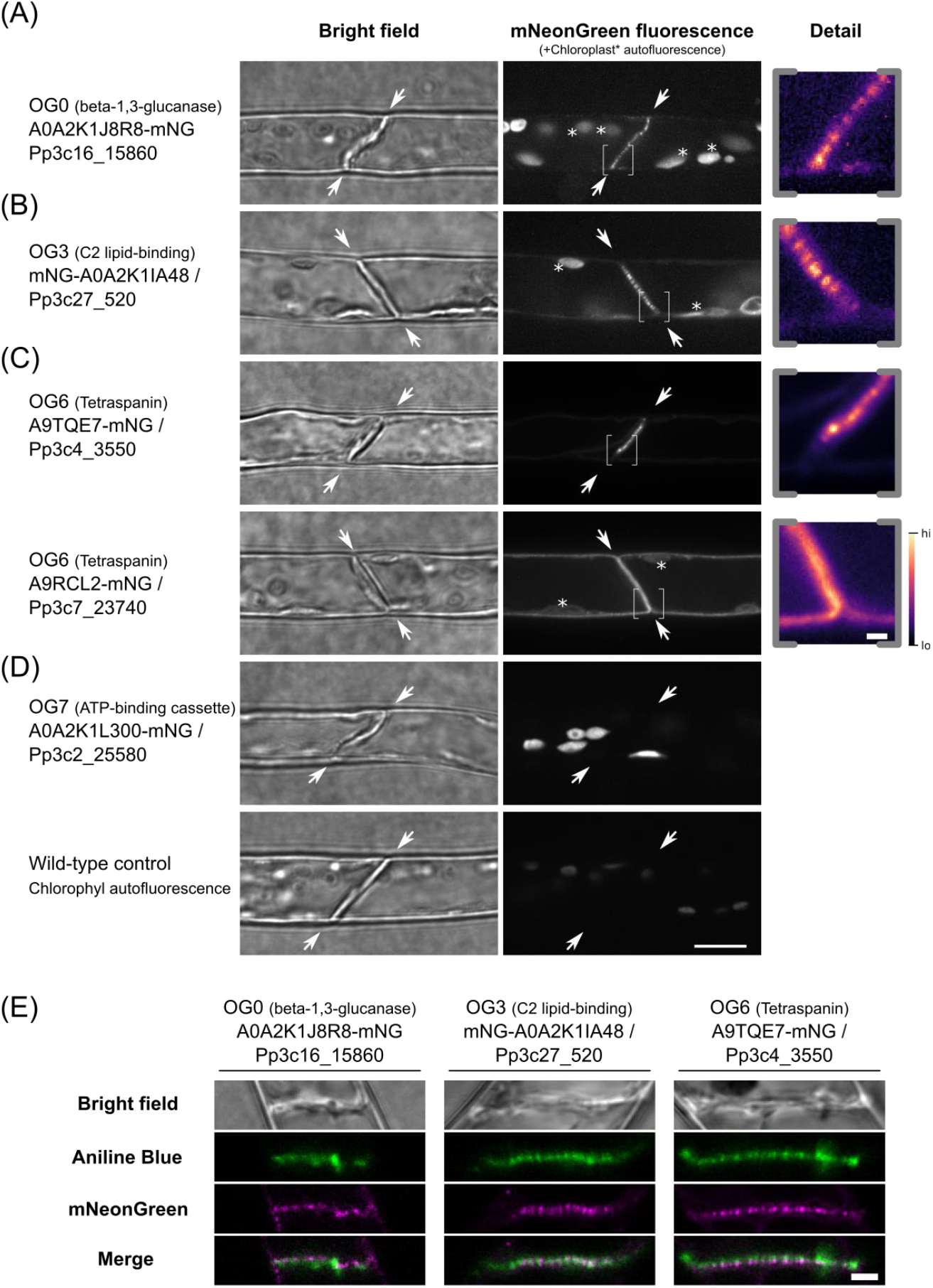
Localization of selected *P. patens* orthogroup members in moss protonemal cells reveals plasmodesmal association. (A-D) Micrographs of moss protonemal cells expressing the indicated protein fused to fluorescent protein mNeonGreen, taken under bright field (left) and fluorescence imaging conditions (right). Proteins belonging to the β-1,3-glucanase (A), C2 lipid-binding (B), Tetraspanin (C) and ATP-binding cassette (D) orthogroups were localized. The dividing interface between two neighbouring cells where plasmodesmata are exclusively located in this tissue are highlighted by arrows. When the fusion protein was detected at this location, an expanded view of part of the dividing wall (indicated by brackets), is shown on the right in pseudocolour. Examples of autofluorescent chloroplasts are marked with an asterisk. The A0A2K1L300-mNG fusion protein localized to chloroplasts as levels of chloroplast autofluorescence under the same imaging and display conditions in wild-type tissue were vastly lower (bottom row). Scale bars are 10 µm in overview images, and 1 µm in expanded views respectively. (E) Co-localization of the three mNeonGreen fusion proteins localizing to the cell interface (Magenta) with callose stain aniline blue (Green). A single confocal plane is depicted showing co-occurrence of the callose and plasmodesmal protein fusion proteins (merge, bottom row). Scale bar is 1 µm.

For OG3, representing the C2 lipid-binding protein family that contains the plasmodesmata-associated MCTPs (Brault et al., 2019), no member was identified in the plasmodesmal fraction from *P. patens* (Table S4). Therefore, we selected A0A2K1IA48 as a candidate *P. patens* plasmodesmal protein as it has the closest homology to Arabidopsis MCTP4 using a phmmer search and is most-abundantly expressed in moss tissues (Ortiz-Ramírez et al., 2016; Fernandez-Pozo et al., 2020). We generated a fluorescent protein fusion by homologous recombination, inserting mNeonGreen at the N-terminus of A0A2K1IA48 and observed a punctate localisation restricted to the cell junction (Fig 2B). Again, aniline blue co-localisation confirmed co-incidence of the signal with plasmodesmal callose, validating A0A2K1IA48 as a plasmodesmal C2 lipid-binding protein from *P. patens*. We also noted that A0A2K1IA48-mNG showed weak ER associated fluorescence that was enriched at discrete foci at the periphery of the external surface of cells (Fig S3), possibly being points of connection between the ER and the plasma membrane as would be expected for proteins in membrane contact sites.

The tetraspanin group OG6 contained 2 members identified in the *P. patens* plasmodesmal fraction: A9RCL2 and A9TQE7. mNeonGreen fusions at the C-terminus of these two tetraspanins revealed two different patterns of localisation. A9RCL2 displayed a punctate pattern of localisation at the cell periphery that co-localised with aniline blue staining of plasmodesmal callose (Fig 2C, E). By contrast, A9TQE7 showed even distribution in the periphery of the cell suggesting it is not enriched in plasmodesmata but present in the entire plasma membrane (Fig 2C). Therefore, we validated only A9RCL2 as a candidate plasmodesmata-associated protein.

OG7 represents ATP-binding cassette proteins that, by contrast with members from OG0, 3 and 6, have never been validated as plasmodesmata-associated proteins in any species. To test whether this group might represent novel core plasmodesmal proteins, we identified A0A2K1L300 in our purified plasmodesmal fraction (Table S2) and inserted mNeonGreen by homologous recombination to generate a A0A2K1L300-mNG fusion. Live imaging shows this protein fusion localizes to chloroplasts suggesting it is not enriched in plasmodesmata (Figure 2D).

### Plasmodesmal association of core orthogroup members is conserved in heterologous species

The validation of plasmodesmal association of *P. patens* proteins from orthogroups represented in plasmodesmal proteomes supports the hypothesis that orthogroup analysis can identify core, conserved plasmodesmal proteins. We reasoned that core, conserved plasmodesmal proteins would be recruited to plasmodesmata in any plant species, *i*.*e*., would localise to plasmodesmata if expressed in a heterologous species. To test this hypothesis, we expressed OG representatives from Arabidopsis and *P. patens* in *Nicotiana benthamiana* leaf epidermal cells and used live cell imaging to determine their association with plasmodesmata. For OG0 we inserted mCitrine downstream of the predicted signal peptide for Q9FHX5 (Arabidopsis BG_PPAP) and mNeonGreen downstream of the catalytic domain of A0A2K1J8R8 and expressed the gene fusions transiently in *N. benthamiana* leaves. Both proteins showed punctate distribution across the cell periphery, with foci of fluorescence co-incident with aniline blue stained plasmodesmal callose (Figure 3A, B).

**Figure 3.**
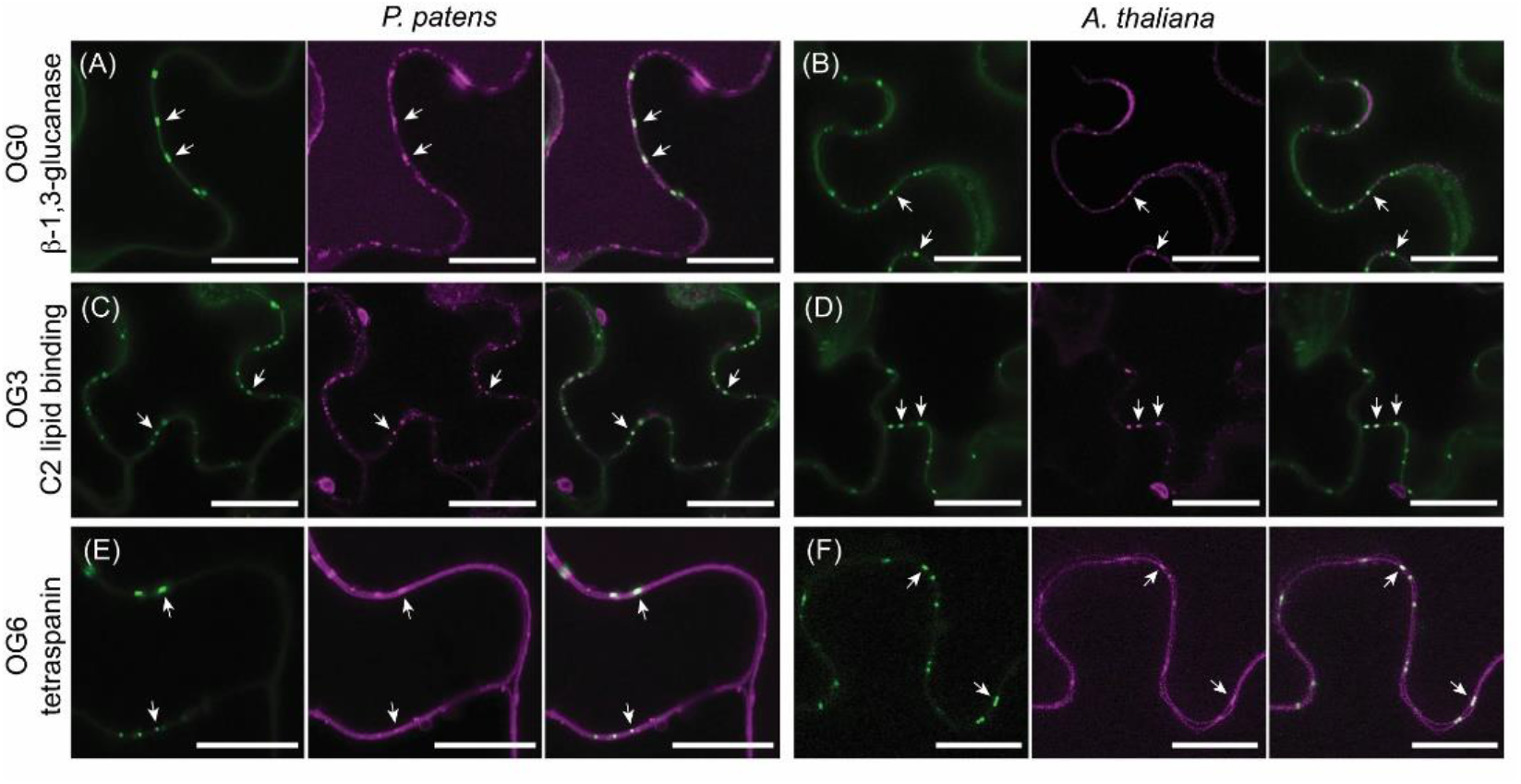
Proteins from OG0, OG3 and OG6 maintain plasmodesmal association in a heterologous species. Confocal micrographs of moss (A, C, E) and Arabidopsis (B, D, F) members of these orthogroups produced in *N. benthamiana* leaf epidermal cells. In each panel, aniline blue stained plasmodesmal callose is green (left), the candidate-FP fusion is magenta (centre) and the overlay is on the right. Members of OG0, representing β-1,3-glucanases, localise to the cell periphery and accumulate at plasmodesmata as indicated by aniline blue co-staining of plasmodesmal callose (arrows). (A) is A0A2KIJ8R8-mNG and accumulates in the vacuole as well as at the cell periphery. (B) is Q9FHX5-Citrine and is detected as diffuse labelling of the cell wall as well as at plasmodesmata. Members of OG3, representing C2 lipid-binding proteins, localise to plasmodesmata accumulate at plasmodesmata as indicated by aniline blue co-staining of plasmodesmal callose (arrows). (C) is A0A2KIA48-GFP and (D) is Q9C8H3-GFP. Members of OG6, representing tetraspanins, localise to the plasma membrane at the cell periphery and accumulate in plasmodesmata as indicated by aniline blue co-staining of plasmodesmal callose (arrows). (E) is A9RCL2-mNG and (F) is Q8S8Q6-mRuby. Scale bars are 20 μm (A, E, F) or 25 μm (B, C, D).

Similarly, we generated C-terminal fusions of OG3 members Q9C8H3 (AtMCTP4) and A0A2K1A48, and OG6 members Q8S8Q6 (AtTET8) and A9RCL2, with GFP or mRuby and observed punctate localisation when expressed in *N. benthamiana* (Figure 3C-F). These punctae co-localised with aniline blue stained callose, confirming these proteins can be recruited to plasmodesmata in a heterologous system. Thus, β -1,3-glucanases (OG0), C2 lipid-binding proteins (OG3) and tetraspanins (OG6) show characteristics of core plasmodesmal proteins.

We further confirmed conservation of plasmodesmal association for C2 lipid-binding proteins and tetraspanins by stable expression of GFP fusions of *P. patens* A0A2K1A48 and A9RCL2 in Arabidopsis. Again, these protein fusions localised in punctae at the cell periphery (Fig S4). Thus, *P. patens* C2 lipid-binding proteins and tetraspanins show conserved association with plasmodesmata in *P. patens, N. benthamiana* and *A. thaliana*.

### Screening of non-membrane proteins for plasmodesmal association in a heterologous system

We observed that conserved plasmodesmal proteins maintain their localisation in heterologous systems. Therefore, we used this approach to test the plasmodesmal association of candidates from orthogroups for which members were not predicted to all have membrane association. We therefore chose Arabidopsis and *P. patens* representatives of OG5 (‘GDSL esterase/lipase’) and OG16 (glycine-rich RNA-binding proteins, GRPs) and screened for plasmodesmal association in *N. benthamiana*. For OG5 we noted that four members were identified in the *P. patens* plasmodesmal fraction. We selected *P. patens* Q4A3V3 (the member identified in plasmodesmal fractions with the highest number of peptide hits) and its closest homologue in Arabidopsis Q9LY84 (AtGELP91) for localisation analysis. C-terminal protein fusions to GFP showed localisation in a cellular reticulum suggestive of the ER (Fig 4A). No clear association with plasmodesmata was detected.

**Figure 4.**
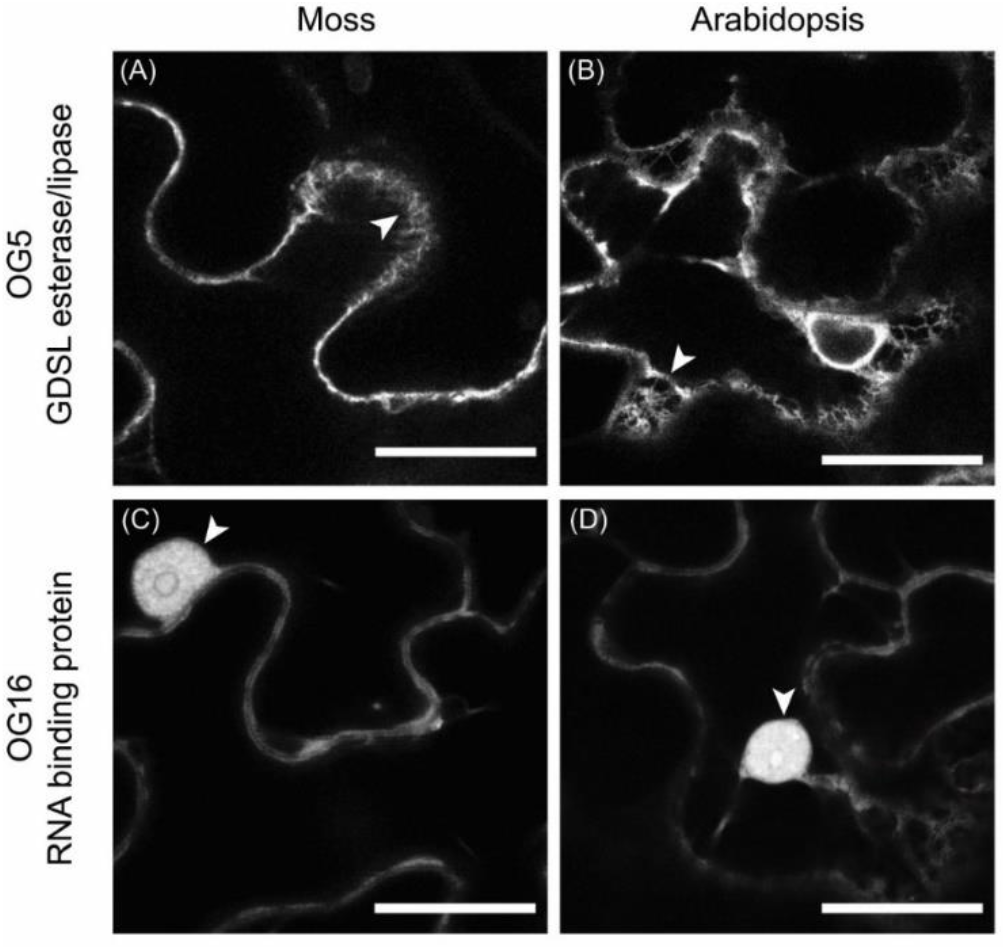
Proteins from OG5 and OG16 don’t accumulate in plasmodesmata. Confocal micrographs of *N. benthamiana* leaf epidermal cells producing fusions of moss (A, C) and Arabidopsis (B, D) members of OG5 and OG16. Members of OG5, containing GDSL esterase/lipases, show an intracellular localisation to a membrane reticulum suggestive of the ER (arrowheads). (A) shows Q4A3V-GFP and (B) shows Q9LY84-GFP. Members of OG16, containing RNA-binding proteins, show nucleo-cytosolic localisation characteristic of soluble proteins. Arrowheads indicate nuclei. (C) shows Q8LPB1-GFP and (D) shows Q03250-GFP. Scale bars are 25 μm.

For OG16 (GRPs), representatives were identified in both Arabidopsis and *P. patens* plasmodesmal proteomes. We selected Q03250 (AtGRP7) from Arabidopsis as it was represented in 2 of 3 Arabidopsis plasmodesmal proteomes, and Q8LPB1 (PpGRP2) from *P. patens* as it had the highest number of unique peptides identified from our *P. patens* fraction. C-terminal fusions of both Arabidopsis and *P. patens* GRPs to GFP showed a nucleo-cytosolic localisation in *N. benthamiana* leaves (Fig 4B). We further tested the localisation of AtGRP7 and PpGRP2 in Arabidopsis by stable transformation and observed similar localisation patterns (Fig S5). Therefore, neither non-membrane associated orthogroup showed plasmodesmal association. Whether this arises because the plasmodesmal fraction of the ER and cytosol cannot be resolved by light microscopy or because these proteins do not associate with plasmodesmata is unclear.

### Phylogenetic analysis within orthogroups to identify different patterns of evolution

Multiple cell components including plasma membrane, cytosol, ER and cell wall, are incorporated into plasmodesmal structure. For some of these compartments such as the plasma membrane and cell wall, plasmodesmal association could plausibly be associated with slow, or even static, protein turnover, allowing detectable accumulation in live imaging approaches. By contrast, cytosolic proteins associated with plasmodesmata might have rapid and transient association with structural components of plasmodesmata or mobile cargoes. While live-imaging can confirm plasmodesmal association of proteins that accumulate at plasmodesmata such that the fluorescence signal associated with plasmodesmata is greater than or separated from the surrounding pool, the approach is limited when accumulation is not a feature of protein behaviour. We failed to confirm plasmodesmal association of OG5 members despite equally strong proteomic support for plasmodesmal association as those of OG6. However, as the GDSL esterase/lipase proteins accumulate in the ER throughout the cell, a discrete association with plasmodesmata might be impossible to resolve by live imaging.

To explore whether protein family phylogenies could identify patterns that might further resolve the likelihood of conserved plasmodesmal association, we generated unrooted cladograms of the protein families than are represented by orthogroup members from Arabidopsis, poplar and moss. We overlayed the resulting trees with proteomic data and assessed whether members identified in plasmodesmal fractions were distributed throughout a tree or were clustered in specific clades that might hint at sub-families with plasmodesmal functions. OG3, OG5 and OG16, all show plasmodesmal association predominantly in a single branch of the tree (Fig 5), suggesting plasmodesmal association and function was gained once during evolution of the protein family. By contrast, plasmodesmal association in OG6 is dispersed across the whole tree suggesting that each tetraspanin ancestor has equal likelihood of being associated with plasmodesmata (Fig 5). OG0 shows no clear phylogenetic pattern associated with plasmodesmal association. As proteins validated as core plasmodesmal proteins are represented amongst trees that harbour single clades and whole tree distribution of proteomic hits, this approach offers no further resolution in identifying core plasmodesmal proteins. However, for protein families with plasmodesmal-association in specific clades, it offers potential to identify candidate plasmodesmal family members from species for which a plasmodesmal proteome has not been generated.

**Figure 5.**
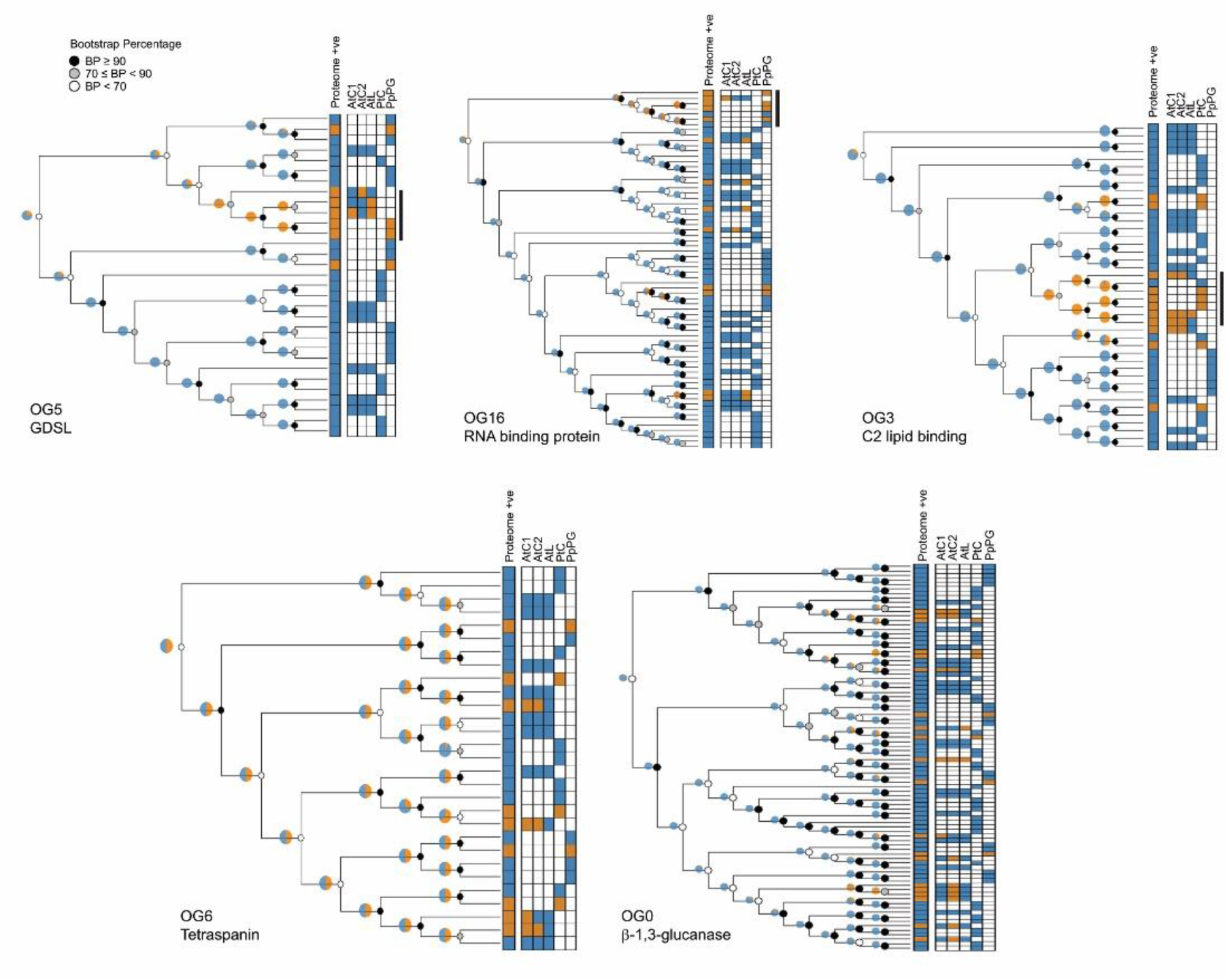
Unrooted cladograms of orthogroup members from *A. thaliana, P. trichocarpa*, and *P. patens* as defined by a hmmsearch with a threshold of E < 1 × 10^−100^ (OG0 / OG3 / OG5) or E < 1 × 10^−50^ (OG6 / OG16). Each tree has a heatmap of proteome matches for each protein in the tree with orange indicating a proteome hit and blue indicating the protein was not detected in the relevant proteome(s). For OG3, OG5, and OG16, plasmodesmal association appears to correlate with a single clade within the tree, indicated by the black bar to the right of the proteome heatmap. Pie charts estimate the likely ancestral plasmodesmal localisation (orange) by phylogenetic backpropagation. Node support is indicated by greyscale circles.

## Discussion

Plasmodesmata are essential features of plant cells but detailed understanding of their structure and function has long been enigmatic. As membrane-rich structures embedded in the cell wall, they can be described as recalcitrant with respect to biochemical extraction and characterisation, and knowledge of their composition has been revealed in a piecemeal fashion despite considerable research efforts. Despite technical challenges, proteomic strategies have underpinned major leaps of understanding in plasmodesmal function and cell-to-cell communication, yielding the primary knowledge that allowed dissection of plasmodesmal responses and dynamics in response to microbe perception and developmental transitions, as well as formulation of the current model for their core structure being a specialised membrane contact site (Tilsner et al., 2016; Sager and Lee, 2018). Recognising the gains to be made by better understanding of the protein composition of plasmodesmata in different tissues and different species we sought to use a phyloproteomic comparison to define a more detailed atlas of plasmodesmal structure and function.

Defining a proteome will always be subject to sampling and technical variation that limits the depth of an analysis of samples from a single technical or biological context as subcellular fractionation and mass spectrometry are inherently noisy techniques (Cargile et al., 2004). The caveat of this is that the most abundant proteins in the preparations will be the most consistently identified, and so some qualitative metric of abundance can be inferred from the repeated presence of a protein. This rationale also applies to a comparison of proteomes of different species in which consistent identification infers conservation and the approach can be used to identify core plasmodesmal proteins; proteins that are identified across multiple species can be considered as core, and thus essential and conserved, components of plasmodesmata. Thus, comparative phylogenetic analysis of proteomes from different species gains power in identifying key plasmodesmal components from inherently noisy datasets.

With the aim of increasing the analytical power of plasmodesmal proteomics, we generated two new plasmodesmal proteomes from differentiated tissues of Arabidopsis and the moss *P. patens*. A phylogenetic approach to comparative analysis of these and existing proteomes from suspension culture cells from Arabidopsis and poplar allowed us to identify protein orthogroups that were represented across samples from different tissues and species, which we hypothesised contain protein classes that are core to plasmodesmal structure and/or function.

Structural components of plasmodesmata such as the ER-derived desmotubule and apoplastic callose have been observed in plasmodesmata across the green lineage (Robards, 1976; Faulkner et al., 2009; Cook et al., 1997; Brecknock et al., 2011; Brunkard and Zambryski, 2017) suggesting that there are conserved, core proteins associated with plasmodesmal structure and function since their first appearance. We reasoned that such core, conserved proteins would likely be associated with plasmodesmata in distantly related plant species and subcellular localisation of proteins from orthogroups containing β-1,3-glucanases (OG0), C2 lipid-binding proteins (OG3) and tetraspanins (OG6) demonstrated this behaviour; verified plasmodesmata-associated representatives from both moss and Arabidopsis were recruited to plasmodesmata in *N. benthamiana*. Despite this evidence of conservation, phylogenetic analysis of the relationships between the moss, poplar and Arabidopsis protein families from which these orthogroups are derived does not reveal a single pattern of evolution. Trees of these protein families show examples in which plasmodesmal association appears to reside in a single clade suggestive of a single gain in an ancestral protein family member and subsequent retention across lineages, while others show a scattered, almost random distribution in our phylogenetic analysis suggestive that protein family members have been independently repeatedly recruited to the plasmodesmal context. Thus, plasmodesmal association of members belonging to different core protein families might be both conserved or repeatedly gained.

Our approach identified and confirmed β-1,3-glucanases and C2 lipid-binding proteins as core and conserved plasmodesmal components. For β-1,3-glucanases this aligns with their characterised role in callose homeostasis at plasmodesmata, with callose deposition detected at plasmodesmata in moss (Fig 2E; Tomoi et al., 2020; Muller et al., 2022) and Arabidopsis (Levy et al., 2007). Further, a β-1,3-glucanase was identified in a plasmodesmal-enriched fraction of *Chara corallina*, suggesting their conservation as a plasmodesmal component spans beyond land plants (Faulkner et al., 2005). Recent structural models of plasmodesmata propose that the ER and plasma membrane in plasmodesmata are a specialised membrane contact site, with C2 lipid-binding domains acting as a critical element that links the two membranes (Tilsner et al., 2016; Brault et al., 2019). As plasmodesmata from moss through to flowering plants have desmotubules, the conservation of C2 lipid-binding proteins in plasmodesmata suggests these proteins are a central element of this structural conservation. We observed that the moss C2 lipid-binding protein A0A2K1A48 localised at plasmodesmata in moss protonema, but also that it showed localisation patterns suggesting it accumulates at other points where the ER sits at the cell periphery (Fig S3). This further supports the likelihood that there is functional conservation between Arabidopsis and moss C2 lipid-biding proteins in membrane contact sites and that membrane contact sites are an ancient feature of plasmodesmal structure.

Tetraspanins also showed conserved localisation across different species but while they have been previously localised to plasmodesmata in Arabidopsis (Fernandez-Calvino et al., 2011) their functional relevance to plasmodesmata is not yet known. Tetraspanins are associated with membrane compartmentalisation, and in animals have been shown to function in the recruitment and activation of signalling components (Kummer et al., 2020; Levy and Shoham, 2005). For tetraspanins, plasmodesmal association is broadly represented across the cladogram (Fig 5). As tetraspanins are found outside plants, across different kingdoms of eukaryotic life, and as our trees are unrooted, it seems unlikely that this suggests tetraspanins were an evolutionary advance that specifically catalysed the formation of plasmodesmata. However, these proteins might be associated with the specialisation of membrane function associated with the evolution of plasmodesmata. Indeed, as plasmodesmal membranes host localised and specialised signalling cascades, tetraspanins might serve to define the plasmodesmal membrane domain and require further investigation.

With callose deposition central to plasmodesmal function we were surprised that our analysis did not detect callose synthases. While this might arise from our fractionation methods being sub-optimal for their extraction, or from using of non-quantitative mass spectrometry methods, we found that if we reduced the stringency of protein identification in both our Arabidopsis leaf and moss plasmodesmal fractions, allowing an identification probability > 50% threshold for peptide and protein identification and a minimum of one sample, we identify an additional 12 orthogroups present in at least 4 of 5 proteomes, one of which contains callose synthases (Tables S1, S2, S5). This low stringency analysis also reveals an orthogroup containing HIPPs, which have been localised to plasmodesmata in Arabidopsis (Guo et al., 2021) and *N. benthamiana* (Cowan et al., 2018), and an orthogroup containing xyloglucan endotransglucosylase/hydrolase which have been confirmed as plasmodesmal proteins in a concomitant comparative proteomic study (Gombos et al., 2022). In essence by requiring the identification of a protein in multiple independent proteomes, we are increasing the *a priori* likelihood of protein identification within a sample. Taking this Bayesian idea, we can reduce the stringent identification criteria of known plasmodesmal proteins, as we are expecting them to appear in the samples. Moreover, proteins which are mis-identified would not be classed as “core”.

In addition to the increased power of analysis by comparative analysis, the data contained herein and in (Gombos et al., 2022) establishes new knowledge of moss plasmodesmata. It has been noted that moss genomes do not encode a family of PDLPs (Vaattovaara et al., 2019), the most studied plasmodesmal proteins to date. As PDLPs positively regulate callose deposition (Cui and Lee, 2016; Caillaud et al., 2014), their absence from moss suggested the possibility that moss plasmodesmata might not be as dynamic, or regulated in the same way, as those in flowering plants (Lee, 2014; Sager and Lee, 2018). However, detection of callose in algae (Faulkner et al., 2009) and bryophytes (Kitagawa et al., 2019; Tomoi et al., 2020; Muller et al., 2022) suggests that callose regulation of plasmodesmata is an ancestral feature. Indeed, as moss plasmodesmata respond to ABA by closing, like the response observed in poplar, the identification of callose degrading enzymes in moss plasmodesmata (Fig 2, Gombos et al., 2022) suggests that this conservation extends to functional regulation of plasmodesmata. This notion is also supported by callose biosynthesis inhibitor treatments in *P. patens* showing an effect on moss shoot branching (Coudert et al., 2015).

Our study further expands the state of knowledge of plasmodesmal proteins to encompass differentiated tissues. Previous proteomes from Arabidopsis and poplar have exploited cell suspension cultures, in which the simple plasmodesmal form is dominant (Bayer et al., 2004). Plasmodesmata increase in complexity with cell expansion and tissue age (Roberts et al., 2001; Faulkner et al., 2008; Fitzgibbon et al., 2013), and transitions in plasmodesmal structure define different functional potential for plasmodesmata (Oparka et al., 1999; Nicolas et al., 2017; Ross-Elliott et al., 2017). Therefore, plasmodesmata of different forms likely recruit specific regulatory machinery that can be uncovered by expanding proteomic knowledge of plasmodesmata from different tissues (Kraner et al., 2017).

Cell-to-cell communication is a central feature of multicellularity and therefore a greater understanding of plasmodesmata promises to greatly enhance our knowledge of a whole range of plant processes by which cells and tissues co-ordinate and communicate to enable organism-level responses. The details of plasmodesmal structure and function are slowly being revealed and we have demonstrated the benefit of enhancing the knowledge gained from technically difficult proteomic profiling by pooling new and existing information to identify conserved, core plasmodesmal components. Indeed, the efforts here and in the Gombos et al. paper offer further opportunity to define the core structure of plasmodesmata and expand our understanding across the evolutionary tree to which future efforts can add mechanistic and physiological understanding.

## Acknowledgements

Work in the Faulkner lab is funded by the European Research Council (grant 725459, “INTERCELLAR”) and the Biotechnology and Biological Research Council Institute Strategic Programme ‘Plant Health’ BBS/E/J/000PR9796. MGJ was funded by a John Innes Foundation Rotation Studentship. We thank Dr. P.A.C. van Gisbergen for assistance with moss callose staining.

## Notes

### Competing Interest Statement

The authors have declared no competing interest.

