## Supplemental figures for "Comparative phyloproteomics identifies conserved plasmodesmal proteins"

### Figure S1

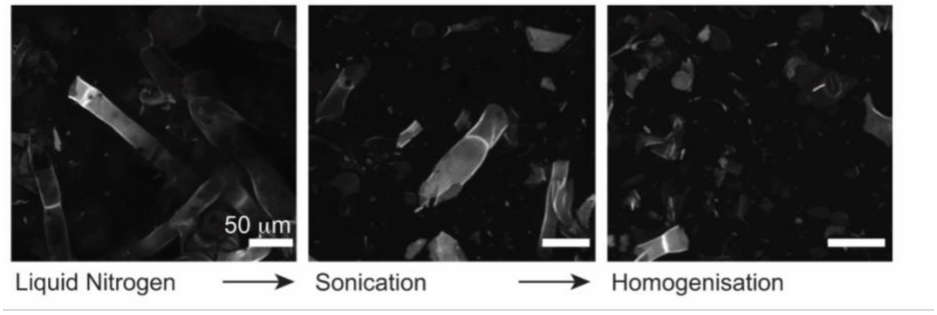


**Figure S1**: Validation of mature plant tissue fractionation of *P. patens* after successive disruption steps. Tissue fragments stained by Calcofluor White observed by confocal microscopy. Scalebar indicates 50 µm.

**Figure S2**

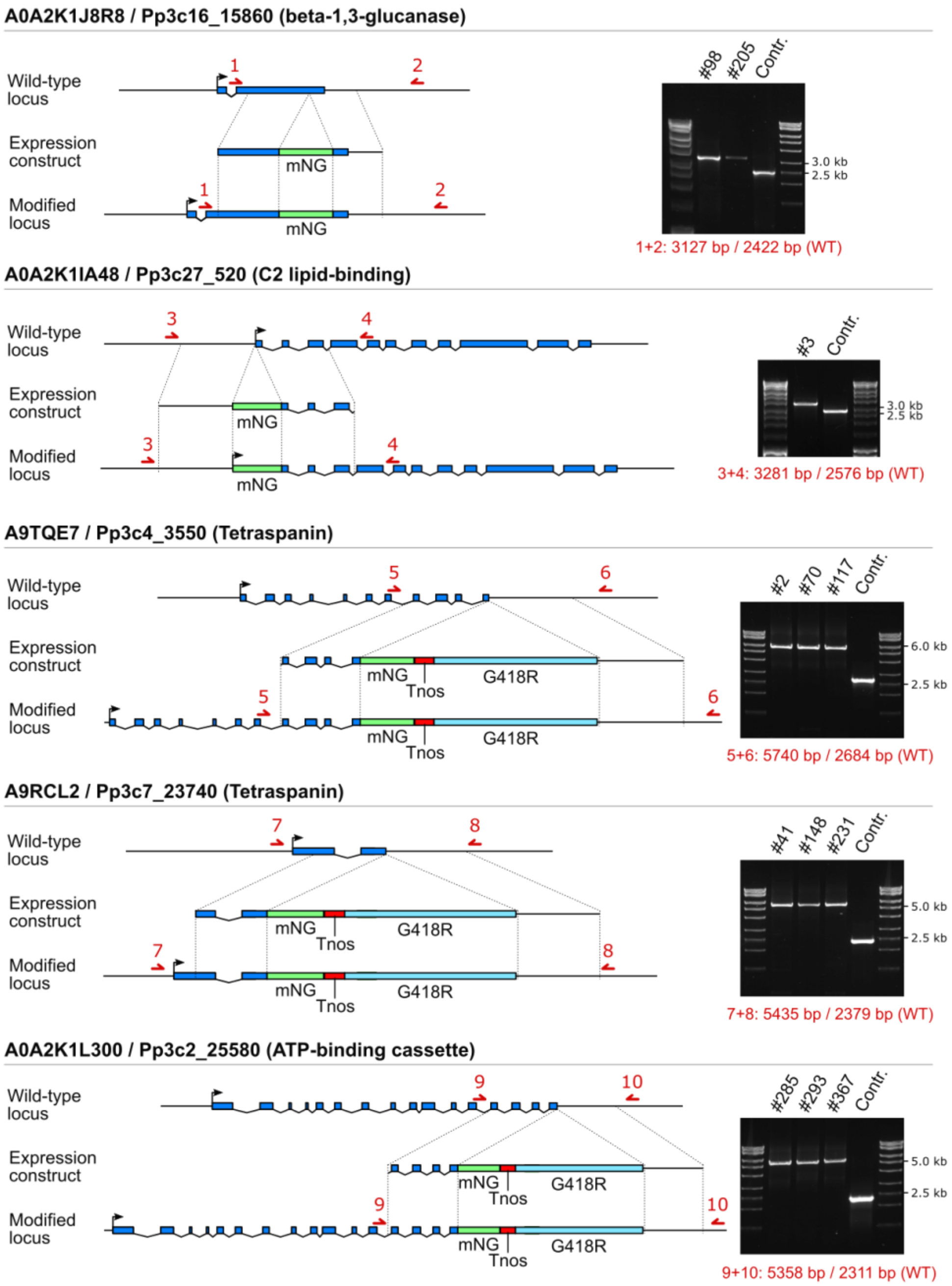


**Figure S2**: Generation and validation of moss strains expressing candidate plasmodesmal proteins fused to fluorescent protein mNeonGreen. Schematic representations of the genomic locus for each indicated gene with its intron-exon structure (blue boxes) and the constructs used for mNeonGreen tagging via homologous recombination (dashed lines) shown. In tagged lines, a fragment containing the mNeonGreen encoding sequence (green box), and in case of C-terminal tagging also a nopaline synthase terminator (red box) and a cassette conferring G418 resistance (light blue box), is integrated at the start, end or within the coding sequence of the native gene. Red arrows and associated numbers denote primer binding sites used for confirmation of the obtained lines by PCR. The products obtained after PCR reaction and their predicted sizes are given on the right. The numbers of analysed transformants are given above the gel images.

**Figure S3**

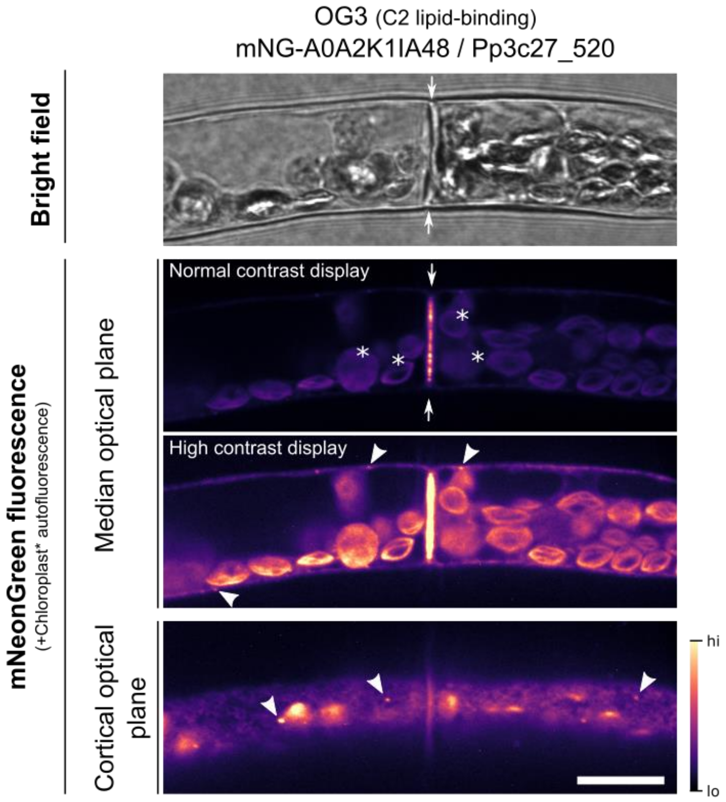


**Figure S3**: Occurrence of labelled C2 lipid-binding protein-positive punctae at the cell periphery in addition to plasmodesmal association in moss. Two protonemal cells expressing mNeonGreen-A0A2K1IA48 are shown under bright field (top) and fluorescence imaging conditions. Confocal slices were acquired both in the median plane of the cells and in the cortical plane (bottom). In addition to strong localization to the cell-cell interface (arrows), under high-contrast display settings distinct punctae at the cell periphery were discernible (arrowheads). In the cortical plane the fusion protein also localizes very weakly to reticulate structures, reminiscent of ER. Scale bar indicates 10 µm.

**Figure S4**

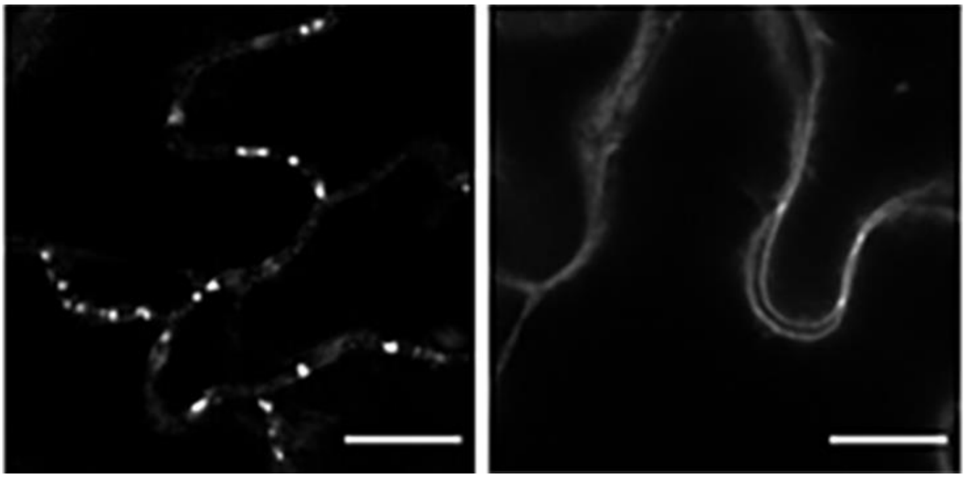

**Figure S4**: Stable expression of the *P. patens* C2 lipid-binding domain protein A0A2K1IA48-GFP (left) and tetraspanin A9RCL2-GFP (right) in Arabidopsis showing accumulation at punctae in the cell periphery suggestive of plasmodesmal association. Scale bar is 10 μm.

**Figure S5**

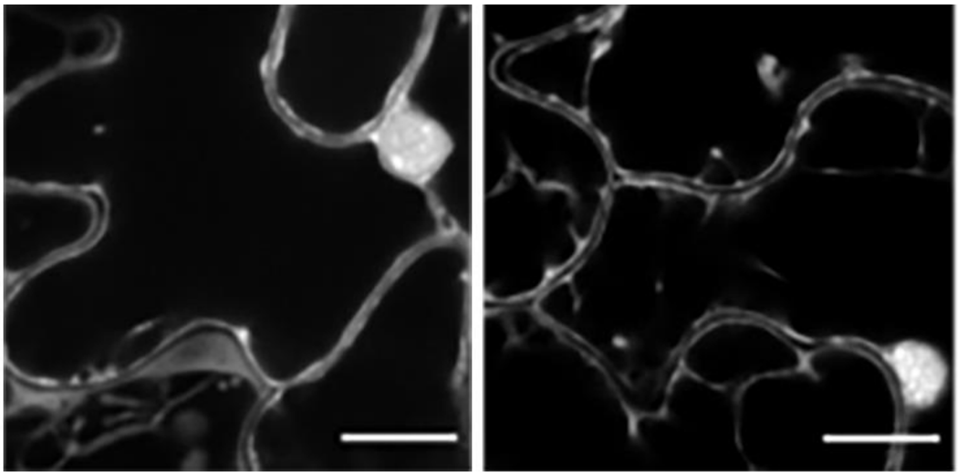

**Figure S5**: Stable expression of the RNA-binding proteins Q8LPB1-GFP (*P. patens*, left) and Q03250-GFP (*A. thaliana*, right) in Arabidopsis showing nucleo-cytosolic localisation. Scale bar is 10 μm.
