## Supplemental Table S5 for "Comparative phyloproteomics identifies conserved plasmodesmal proteins"

| Orthogroup | Protein Class | # Proteomes | # Proteins | In Table 1 |
|---|---|---|---|---|
| OG0 | β-1,3-glucanase | 5 | 14 | Yes |
| OG1 | Peroxidase | 4 | 25 | Yes |
| OG2 | Lectin receptor-like kinase | 5 | 56 | No |
| OG3 | C2 lipid-binding | 4 | 20 | Yes |
| OG4 | SKU5 | 4 | 14 | Yes |
| OG5 | GDSL esterase/lipase | 4 | 17 | Yes |
| OG6 | Tetraspanin | 4 | 5 | Yes |
| OG7 | ATP-binding cassette | 5 | 11 | Yes |
| OG8 | Aspartyl protease | 4 | 10 | Yes |
| OG9 | Leucine-rich repeat receptor-like kinase | 4 | 10 | Yes |
| OG10 | Leucine-rich repeat extensin-like | 4 | 12 | Yes |
| OG11 | Callose synthase | 4 | 15 | No |
| OG12 | NDR1/HIN1-like protein | 5 | 10 | No |
| OG13 | Histone H2B | 4 | 9 | Yes |
| OG14 | Tubulin beta-7 | 4 | 14 | Yes |
| OG15 | Transmembrane protein | 5 | 10 | No |
| OG16 | Glycine-rich RNA-binding | 4 | 13 | Yes |
| OG19 | DUF26 containing protein | 4 | 8 | Yes |
| OG23 | Heavy metal associated isoprenylated plant protein | 4 | 9 | No |
| OG28 | Eukaryotic initiation factor 4A | 4 | 7 | Yes |
| OG33 | CSC1-like protein ERD4 | 5 | 7 | No |
| OG37 | Xyloglucan endotransglucosylase/ hydrolase | 4 | 7 | No |
| OG40 | Subtilisin-like protease | 4 | 6 | Yes |
| OG45 | Serine carboxypeptidase-like | 4 | 6 | No |
| OG50 | Calcium-dependent lipid-binding | 5 | 5 | Yes |
| OG63 | Ribosomal protein | 4 | 4 | Yes |
| OG71 | Polyubiquitin | 4 | 5 | No |
| OG73 | Prohibitin | 4 | 5 | No |
| OG94 | Transmembrane protein | 4 | 4 | No |
| OG107 | β-glucosidase | 4 | 4 | No |
